# Genome-wide association study implicates *CHRNA2* in cannabis use disorder

**DOI:** 10.1101/237321

**Authors:** Ditte Demontis, Veera Manikandan Rajagopal, Thomas D. Als, Jakob Grove, Jonatan Pallesen, Carsten Hjorthøj, Per Qvist, Jane Hvarregaard Christensen, Jonas Bybjerg-Grauholm, Marie Bækvad-Hansen, Laura M. Huckins, Eli A. Stahl, Allan Timmermann, Esben Agerbo, David M. Hougaard, Thomas Werge, Ole Mors, Preben Bo Mortensen, Merete Nordentoft, Mark Daly, Mette Nyegaard, Anders D. Børglum

## Abstract

Cannabis is the most frequently used illicit psychoactive substance worldwide^1^. Life time use has been reported among 35-40% of adults in Denmark^2^ and the United States^3^. Cannabis use is increasing in the population^4–6^ and among users around 9% become dependent^7^. The genetic risk component is high with heritability estimates of 51^8^–70%^9^. Here we report the first genome-wide significant risk locus for cannabis use disorder (CUD, P=9.31×10^−12^) that replicates in an independent population (P_replication_=3.27×10^−3^, P_metaanalysis_=9.09×10^−12^). The finding is based on a genome-wide association study (GWAS) of 2,387 cases and 48,985 controls followed by replication in 5,501 cases and 301,041 controls. The index SNP (rs56372821) is a strong eQTL for *CHRNA2* and analyses of the genetic regulated gene expressions identified significant association of *CHRNA2* expression in cerebellum with CUD. This indicates a potential therapeutic use in CUD of compounds with agonistic effect on the neuronal acetylcholine receptor alpha-2 subunit encoded by *CHRNA2*. At the polygenic level analyses revealed a significant decrease in the risk of CUD with increased load of variants associated with cognitive performance.

## Main

Overall the prevalence of diagnosed CUD in the population has been estimated to 1-1.5% among Europeans^10,11^ and Americans^5^. CUD is associated with a range of adverse health problems^12,13^ including risk of psychosis^14,15^, bipolar disorder^16^, anxiety disorder^17^ and cognitive impairment with more persistent use associated with greater decline^18^. Estimates of the heritability for cannabis use initiation and life-time cannabis use, with respect to the amount of variance explained by common variants (i.e. the SNP heritability), has been estimated to 0.06^19^ - 0.2^20^. Four GWASs related to cannabis use have been conducted without genome-wide significant findings: one study of cannabis dependence^21^ and three studies of lifetime cannabis use^19,20,22^. In addition, two recent GWASs have reported genome-wide significant associations, albeit with negative or ambiguous replication results: a study of DSM-IV cannabis dependence criterion counts in a combined sample of 14,754 European Americans and African Americans^23^, which reported three genome-wide significant loci associated with cannabis use severity; and a GWAS of cannabis dependence of 2,080 European cases and 6,435 controls which identified one genome-wide significant locus^24^.

Here we present results from a GWAS and subsequent replication based on analyses of the largest cohorts of diagnosed CUD reported so far. Individuals included in the discovery GWAS come from the Danish nation-wide population based cohort collected by the Lundbeck Foundation Initiative for Integrative Psychiatric Research (iPSYCH)^25^. The iPSYCH cohort was ascertained to study six major psychiatric disorders (schizophrenia, bipolar disorder, major depressive disorder, attention-deficit hyperactivity disorder (ADHD), anorexia nervosa and autism spectrum disorder) and consists of 79,492 genotyped individuals. The present GWAS was based on 2,387 individuals with a diagnosis of CUD (ICD10 F12.1-12.9) and 48,985 individuals not diagnosed with CUD, all identified in the iPSYCH cohort. Data analysis was conducted using the Ricopili pipeline^26^, including stringent quality control of genotyped variants and individuals (online methods). Information about non-genotyped markers was obtained by imputation^27,28^ using the 1000 genomes phase 3 as reference panel^29^. GWAS was performed using imputed marker dosages and an additive logistic regression model with relevant principal components to correct for confounders such as population stratification, and the major psychiatric disorders studied in iPSYCH as covariates. Only markers with an imputation INFO score > 0.7, minor allele frequency (maf) > 0.01 and bi-allelic markers were retained, in total 8,971,679 genetic markers. We identified 26 genome-wide significant SNPs (P< 5×10^−8^), located in a single locus on chromosome 8 (Figure 1 and 2). The index SNP, rs56372821, showed an odds ratio of 0.73 (P= 9.31×10^−12^) with respect to the minor allele A (Table 1, Figure 1 and 2).

**Figure 1.**
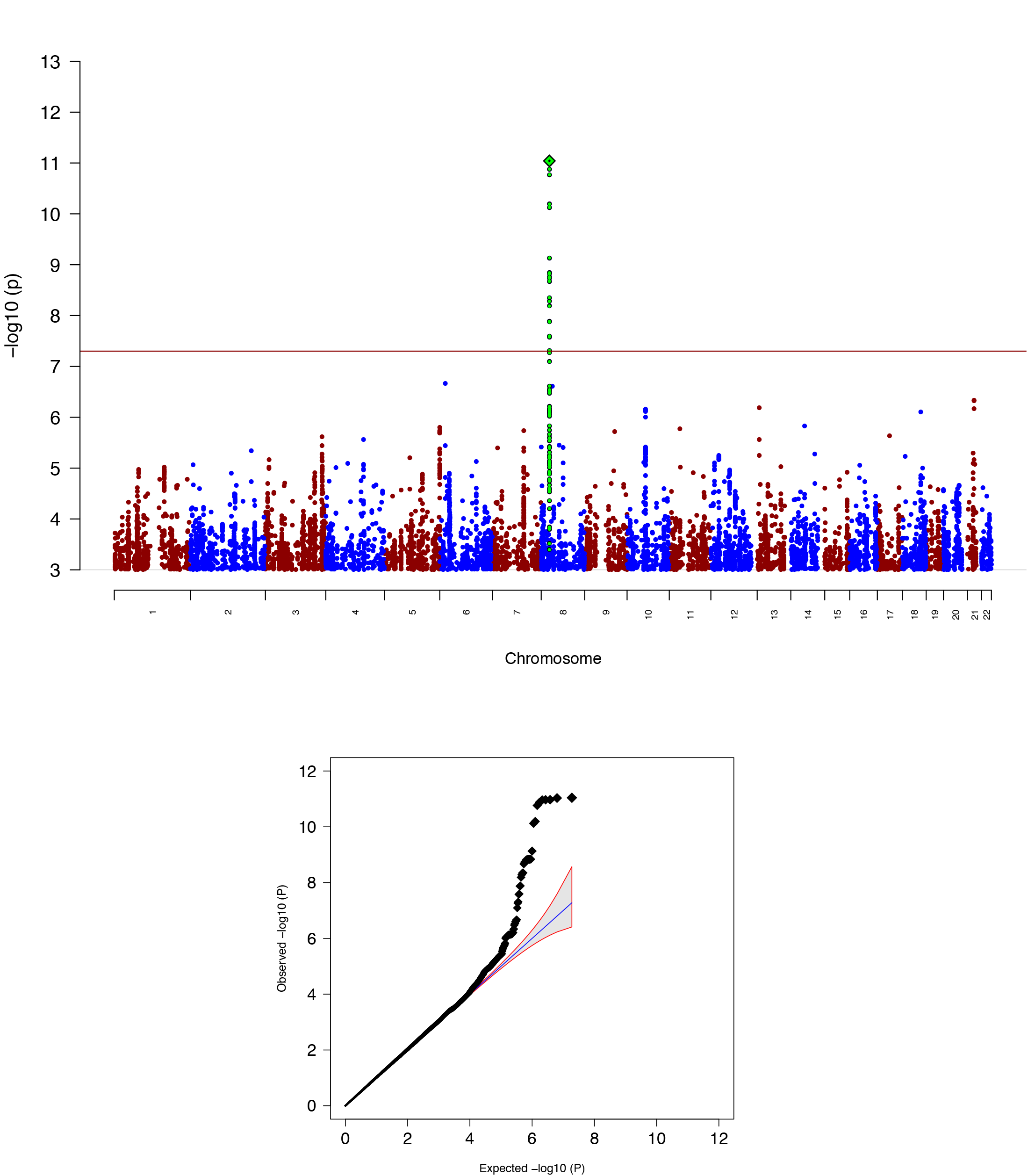
**A)** Manhattan plot Manhattan plot of the results from the GWAS of CUD. The index is highlighted as a green diamond and SNPs in high LD with the index SNP are marked in green. **B)** Quantile-quantile plot of the expected and observed P-values from GWAS of CUD.

**Figure 2.**
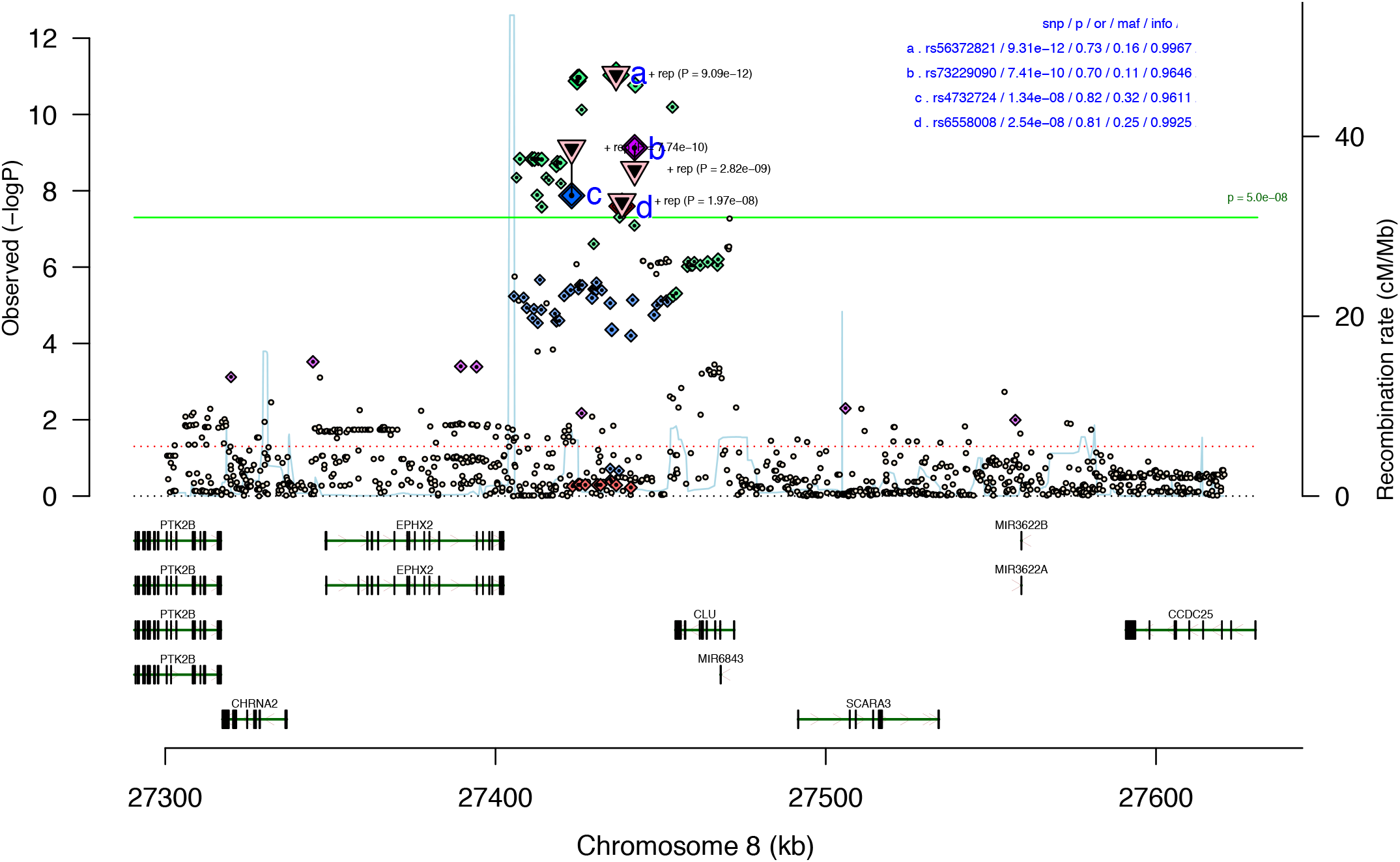
Regional association plot of the local association results for the risk locus at chromosome 8. The index SNP (rs56372821) and additional three correlated genome-wide significant SNPs (LD with index SNP: 0.2 < r^2^ < 0.7) are marked with letters (a-d), the triangle represents the P-value from metaanalysis with the replication cohort from deCODE. The location and orientation of the genes in the region and the local estimates of recombination rate is shown. The association P-value (p), odds ratio (or), minor allele frequency (maf) and imputation info-score (info) is presented in upper right corner.

**Table 1.** Genome-wide significant SNPs associated with cannabis use disorder. Results for genome-wide significant SNPs in the associated locus at chromosome 8 in the primary GWAS and/or in the meta-analysis with results from an independent cohort collected by deCODE. Results for the index SNP (Index SNP) is shown together with results from supporting correlated SNPs in the locus (0.2 < r^2^ < 0.7) (LD SNPs). Alleles for the variants (A1 and A2), frequency of A1 in controls (Frq A1 con), the odds ratio (OR) for the effect of A1, and P-values (P) are given.

| Index SNP | LD SNPs | CHR | BP (hg19) | iPSYCH |  |  |  |  | deCODE |  |  |  |  | meta-analysis |  |  |  |
| --- | --- | --- | --- | --- | --- | --- | --- | --- | --- | --- | --- | --- | --- | --- | --- | --- | --- |
|  |  |  |  | A1 | A2 | Frq A1 con | OR | P | A1 | A2 | Frq A1 con | OR | P | A1 | A2 | OR | P |
| rs56372821 |  | 8 | 27436500 | A | G | 0.163 | 0.728 | **9.31x10<sup>-12</sup>** | A | G | 0.144 | 0.878 | 3.27 x10<sup>-03</sup> | A | G | 0.803 | **9.09 x10<sup>-12</sup>** |
|  | rs4732724 | 8 | 27423062 | C | G | 0.324 | 0.820 | **1.34 x10<sup>-08</sup>** | C | G | 0.367 | 0.905 | 1.60 x10<sup>-03</sup> | C | G | 0.866 | **7.74 x10<sup>-10</sup>** |
|  | rs6558008 | 8 | 27438306 | A | C | 0.755 | 1.237 | **2.54 x10<sup>-08</sup>** | C | A | 0.228 | 0.915 | 1.45 x10<sup>-02</sup> | A | C | 1.159 | **1.97 x10<sup>-08</sup>** |
|  | rs73229090 | 8 | 27442127 | A | C | 0.110 | 0.702 | **7.41 x10<sup>-10</sup>** | A | C | 0.105 | 0.881 | 1.30 x10<sup>-02</sup> | A | C | 0.797 | **2.82 x10<sup>-09</sup>** |
|  | rs937220 | 8 | 27429730 | A | G | 0.259 | 0.8258 | 2.46 x10<sup>-07</sup> | A | G | 0.269 | 0.912 | 7.56 x10<sup>-03</sup> | A | G | 0.871 | **4.53 x10<sup>-08</sup>** |
|  | rs17466060 | 8 | 27422740 | A | G | 0.418 | 0.8594 | 3.94 x10<sup>-06</sup> | A | G | 0.430 | 0.900 | 5.11 x10<sup>-04</sup> | A | G | 0.881 | **1.29 x10<sup>-08</sup>** |

The genome-wide significant locus on chromosome 8 was replicated in an independent European sample consisting of 5,501 cases with CUD and 301,041 population controls (deCODE cohort). The cases were diagnosed with CUD while undergoing inpatient treatment at the SAA Treatment Centre, Vogur Hospital in Iceland (www.saa.is). We tested nine markers located in the risk locus in the Icelandic sample; the index SNP and eight correlated variants (0.2 < r^2^ < 0.7) with P values less than 1×10^−6^ (four genome-wide significant). All variants demonstrated consistent direction of association and the most strongly associated variant (rs56372821) in the discovery GWAS had a P-value of 3.27×10^−3^ in the deCODE cohort. In the meta-analysis rs56372821, became slightly stronger associated with CUD (P = 9.09×10^−12^), and additional two variants became genome-wide significant (Table 1).

There was no evidence of association of previously identified genome-wide significant cannabis risk varaints with CUD in our analyses^23,24^ (Supplementary Table 1).). This might be due to different phenotype definitions among the studies as Sherva et al.^23^ analysed association with cannabis criterion counts, and Agrawal et al.^24^ used cannabis exposed (but not dependent) individuals as controls in their study. Additionally, the composition of the cohorts analysed also differ as the previous GWASs were based on cohorts established to study genetics of substance use disorders while the iPSYCH cohort is ascertained for major mental illnesses (Supplementary Table 2).

To assess the proportion of phenotypic variance explained by common variants we applied LD score regression^30^ and the GREML method implemented in GCTA^31^, assuming a population prevalence of 1% for cannabis use disorder (Online methods). Estimates of the liability-scale SNP heritability were h^2^_snp_= 0.09 (SE=0.03) and h^2^_snp_= 0.042 (SE=0.014) using LD score regression and GCTA, respectively. The results imply a smaller contribution from common variants than was previously found for cannabis use^20^. The estimate would probably increase with larger sample size, as a result of a decrease in the size of the error terms of the SNP effect estimates^32^.

We found no signs of contributions from confounding factors like population stratification and cryptic relatedness to the inflation in the distribution of the test statistics (see quantile-quantile plot, Figure 1.B.) using LD score regression, since the intercept was one (intercept = 0.996; SE=0.0079) (online methods).

In the GWAS we corrected for diagnoses of the major psychiatric disorders studied in iPSYCH (see Supplementary Table 2 for distribution of psychiatric disorders among CUD cases and the control group). In addition, we evaluated the impact on the odds ratio for the index SNP by leave-one-out analyses excluding psychiatric phenotypes one at a time in the association analysis (online methods). The odds ratio remained stable (Table 2), supporting that the association was independent of a diagnosis with one the psychiatric disorders evaluated.

**Table 2.**
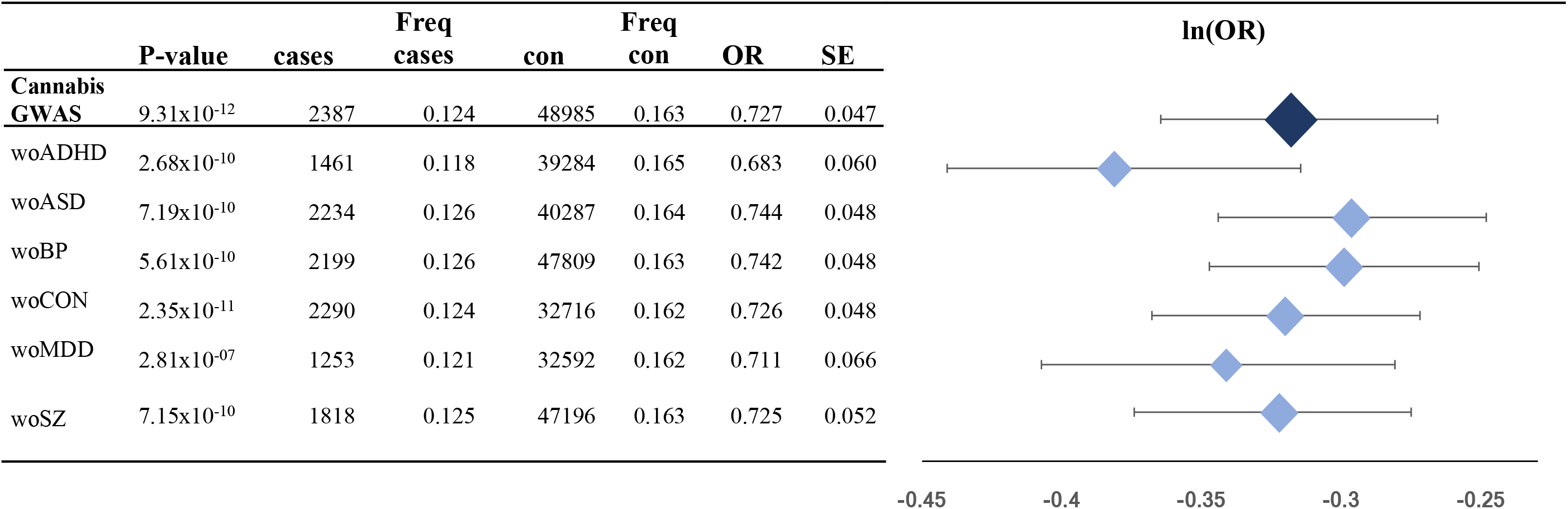
The effect on the odds ratio of rs56372821 of leave-one-phenotype out association analysis. In the analyses individuals with a psychiatric diagnosis are excluded one disorder at a time (without ADHD (woADHD), without autism spectrum disorder (woASD), without bipolar disorder (woBP), without the iPSYCH control cohort (woCON), without major depressive disorder (woMDD), without schizophrenia (woSZ). The P-value for association with CUD (P-value), number of cases (cases), frequency of the minor allele in cases (Freq cases), number of controls (con), frequency of the minor allele in controls (Freq con), odds ratio (OR) and standard error (SE), are given. For illustration ln(OR), and the corresponding standard error.

The risk locus on chromosome 8 has also been found to be genome-wide significantly associated with schizophrenia^26^. In our leave-one-phenotype out analysis the locus remained genome-wide significant when individuals with schizophrenia were removed, excluding any potential confounding from schizophrenia (Table 2). The signal observed for the index SNP (rs73229090) in the GWAS meta-analysis of schizophrenia^26^ is consistent with the direction of association observed in our analysis. Since individuals with schizophrenia often use cannabis (around 13% - 16%^33,34^) it could be speculated that the significant signal observed in schizophrenia, is driven by a sub-group of schizophrenia cases also having CUD. This hypothesis is supported by analysis of the association of rs56372821 with schizophrenia in the iPSYCH sample. We found a nominal significant association of the SNP with schizophrenia when including individulas with co-morbid CUD (2,281 schizophrenia cases and 23,134 controls; OR=0.9; P=0.036), while after exclusion of individuals with CUD (556 cases and 101 controls excluded) the association signal disappeared (1,727 cases, 23,033 controls; OR = 0.97, P=0.63). In order to evaluate further the impact of CUD comorbidity on the odds ratio of rs56372821 in schizophrenia a null distribution for the odds ratio was generated by performing 10,000 permutations randomly removing 556 and 101 individuals among the cases and controls, respectively (Supplementary Figure 1). The observed odds ratio for rs56372821, when excluding individuals with comorbid CUD was 0.97, and differed significantly from random removal of the same number of cases and controls (P_two-sided_=0.0027, P_one-sided_=0.0015). Thus the permutation test supports the hypothesis that the subgroup diagnosed with CUD among schizophrenia cases drives the nominal association observed for rs56372821 in the iPSYCH schizophrenia sample.

We performed a CUD-only analysis testing all SNPs in LD with the index SNP (19 SNPs; r^2^ > 0.7) for association with age at first diagnosis. This analysis suggested the risk alleles to be associated with earlier age at first diagnosis (most significant SNP: rs35236974, P = 0.020). On average CUD cases homozygous for the protective allele got their diagnosis (mean age: 22.41; st.dev = 3.63) one year later than CUD cases having at least one risk allele (mean age: 21.01; st.dev = 3.56).

Among the brain tissues analysed in GTEx the index SNP rs56372821 was found to be a strong eQTL for *CHRNA2* in cerebellum (P-value in GTEx =2.1×10^−7^), with the risk allele (G-allele) being associated with decreased expression of the gene (Supplementary Figure 2). In order to further evaluate the potential regulatory impact of the identified locus on chromosome 8 as well as gene expression differences between cases and controls genome-wide, we imputed the genetically regulated gene expression in 11 brain tissues using PrediXcan^35^. The SNP weights used were derived from models trained on reference transcriptome data sets including 10 brain tissues from GTEx^36^ and transcriptome data from dorsolateral prefrontal cortex generated by the CommonMind Consortium^37,(Huckins et al. submitted to Nature Genetics)^. We tested for association of expression of 2,460 – 10,930 protein coding genes (depending on the tissue; see Supplementary Table 3) with CUD using logistic regression corrected by the same covariates as in the GWAS. One gene, *CHRNA2*, was significantly differently expressed between cases and controls (P = 2.713×10^−6^; beta = −0.21; SE = 0.045) in the cerebellum. The expression model for *CHRNA2* in cerebellum was based on 47 SNPs including four genome-wide significant SNPs rs59724122, rs73229093, rs7838316 and rs11783093 (Figure 4). *CHRNA2* expression was predicted with a valid model in two additional two brain tissues with nominal significant under-expression in cases compared to controls (dorsolateral prefrontal cortex, P=5.19×10^−4^; cerebellar hemisphere P=5.30×10^−3^). That the risk locus for CUD can be linked to *CHRNA2* expression is also supported by Won et al.^38^ who generated high-resolution 3D maps of chromatin contacts (by Hi-C sequencing approach), in order to capture the functional relationships between regulatory and transcribed elements during human corticogenesis. They applied their map to the set of credible SNPs from the large PGC schizophrenia GWAS^26^ and found physical interaction of the genome-wide significant risk locus on chromosome 8, with the regulatory region of *CHRNA2*.

**Figure 3.**
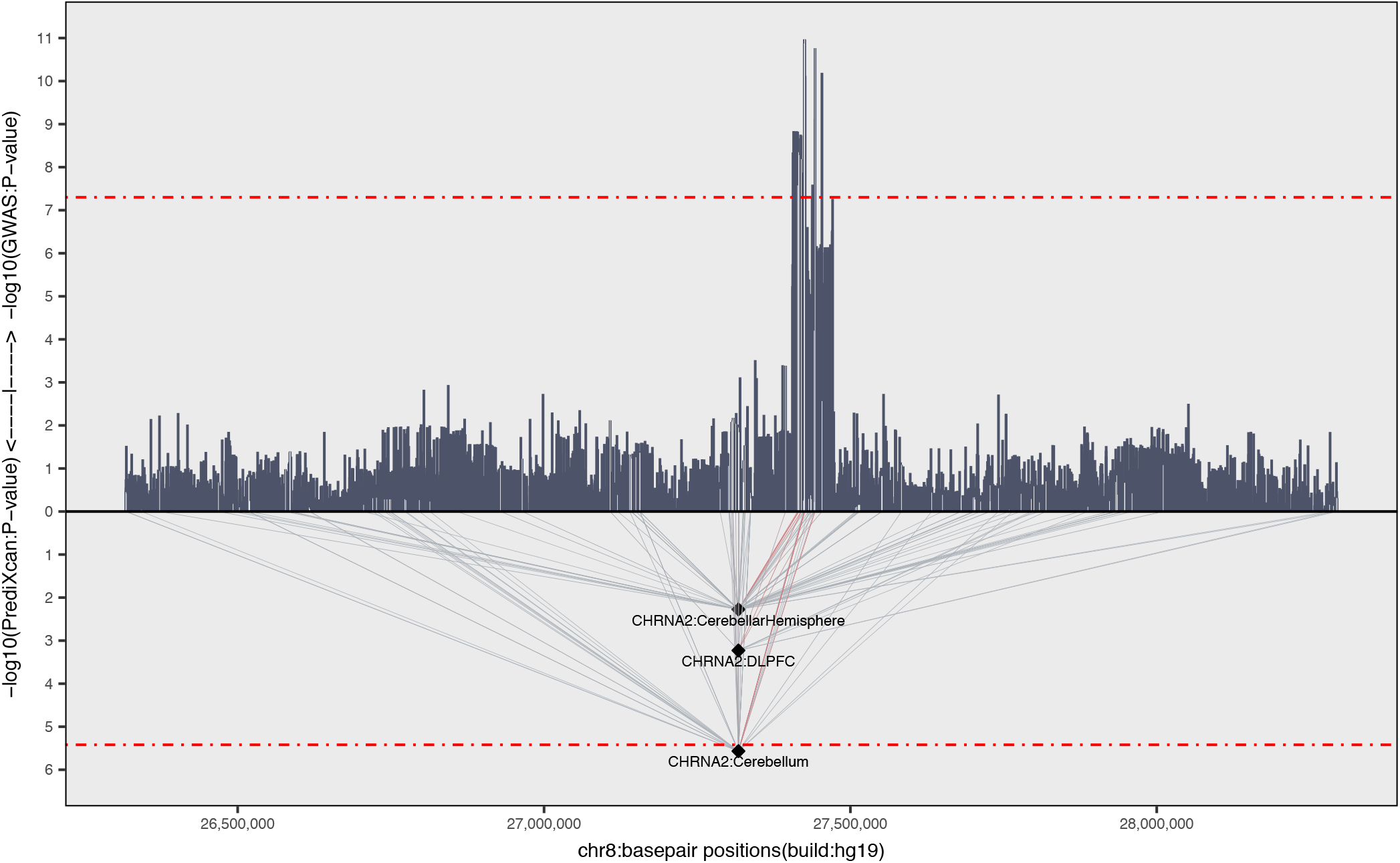
Association of the imputed genetically regulated expression of *CHRNA2* in three brain tissues with a valid model (cerebellar hemisphere, dorsolateret prefrontal contex and cerebellum). The P-value for association of expression with CUD (-log10(PrediXcan:P-value) and P-value from CUD GWAS (−log10(GWAS:P-value)) is given on the y-axis, with a red dotted line indicating statistical significance. Chromosome position is given on the x-axis and the light grey lines indicate which SNPs that are included in the models used to predict gene expression. The thin red lines indicate genetic predictors that are genome-wide significantly associated with CUD.

**Figure 4.**
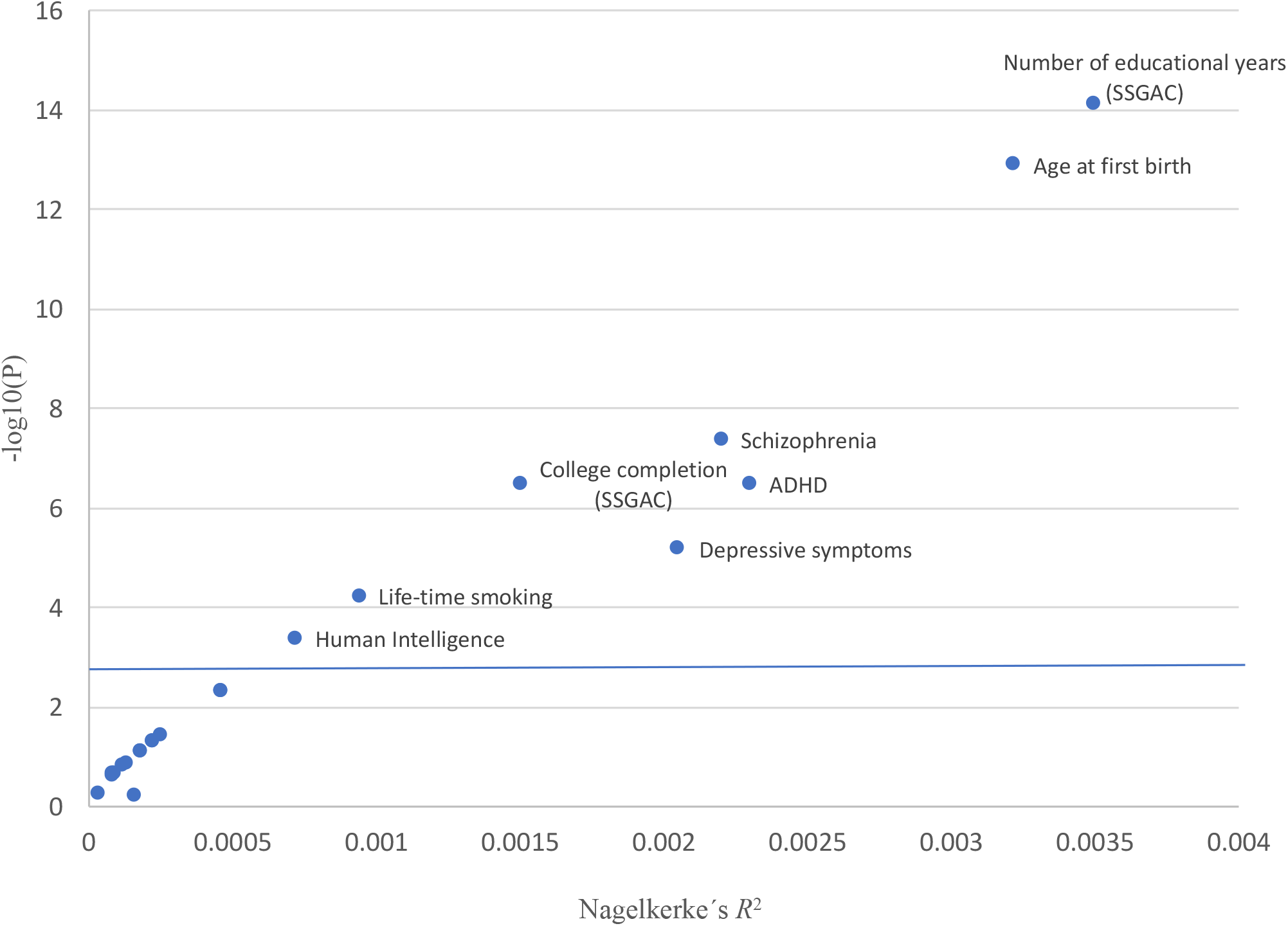
Association of PRS with CUD. PRSs was generated for phenotypes related to cognition, personality, psychiatric disorders, reproduction and smoking behavior based on summary statistics from 22 published GWASs. The variance explained by the scores (Nagelkerke-*R*^2^) is given on the x-axis and the P-value for association of the PRS with CUD on the y-axis.

*CHRNA2* is expressed in the brain^39,40^ and encodes the neuronal acetylcholine receptor (nAChR) alpha-2 subunit, which is incorporated in heteropentameric neuronal nAChRs mainly with the beta-2 or beta-4 subunits^41,42^. Candidate gene studies of common variants have linked this gene to e.g. substance abuse^43^ and nicotine dependence^44,45^ but no genome-wide significant findings have connected *CHRNA2* to any substance abuse or psychiatric disorder, besides the fuctional link between schizophrenia risk variants and *CHRNA2* expression identified by Won et al^38^ described above.

The result suggests *CHRNA2* underexpression in cerebellum (and potentially other brain regions, Supplementary Table 3), to be involved in CUD. Cerebellum may play a role in addiction with respect to reward and inhibitory control^46,47^, and reward-anticipating responses^48^. The cerebellum has high density of cannabinoid receptor 1 (CB1), which mediates the effect of delta9-tetrahydrohydrocannabinol (THC)^49^, the psychoactive compound in cannabis and has also been found to be affected by cannabis use in neuroanatomic studies^50–53^.

There is no reported functional link between cannabis use and alpha-2 subunit containing nAChRs. We hypothesize three potential ways of the involvement of *CHRNA2* in CUD: 1) Substances in cannabis might interact directly with the alpha-2 subunit containing nAChRs as studies have found cannabidiol, a non-psychoactive component of cannabis, to inhibit the alpha-7 containing nAChRs^54^. 2) Cannabis could indirectly affect the alpha-2 subunit containing nAChRs. After binding of an agonist e.g. acetylcholine, the nAChR responds by opening of an ion-conducting channel across the plasma membrane. This causes depolarization of the membrane and can result in presynaptic neurotransmitter release^55^ including dopamine^56,57^. Since the psychoactive active compound of cannabis THC has been found to affect the release of acetylcholine in various brain regions^58,59,60^ it could be speculated that this, through alpha-2 subunit containing nAChRs, could affect dopamine release, a known neurotransmitter involved in addiction^61^. 3) There could be a strong biological link between expression of *CHRNA2* and the cannabinoid receptor 1 gene (*CNR1*). This hypothesis is based on evaluation of gene-expression correlations from genome-wide microarray gene expression profiles in the Allan Brain Atlas^62^ (<u>http://www.brain-map.org/</u>). We found that of all genes evaluated (58692 probes analyzed), *CNR1* demonstrated the strongest negative correlation with *CHRNA2* expression (r_max_= −0.498; Supplementary Figure 3). The signal was driven by opposite expression patterns in a large number of brain tissues, e.g in cerebellum where *CNR1* had a relatively high expression in the cerebellar cortex and *CHRNA2* had a relatively low expression and the opposite was observed for cerebellar nuclei (Supplementary Figure 3). This suggests the existence of a currently uncharacterized biological interaction between the endocannabinoid system and alpha-2 subunit containing nAChR, by which the identified risk locus which is associated with decreased *CHRNA2* expression could be related to increased *CNR1* expression.

The observed association of *CHRNA2* unerexpression with CUD, also implies that the alpha-2 subunit can be a potential drug target in the treatment of CUD, by the use of an agonist selective for alpha-2 subunit containing nAChRs^63,64^. The impact of some compounds with an agonistic effect on the alpha-2 subunit containing nAChRs, like NS9283^65^, ABT-418^66^ and TC-1734^67^, have already been studied primarily due to their potential impact on memory and cognitive processes^65,67–69^. Additionally TC-1734 has been tested in clinical phase I and II trials for its improvement on memory^67,69,70^, and the favorable safty profile of this drug suggests a potential re-purposing for CUD treatment.

*CHRNA2* is related to other nicotine receptor genes that have been identified as risk loci for nicotine dependence and smoking behavior^71,72^. It is therefore relevant to question if our finding is confounded by smoking. In the iPSYCH sample^25^ there is no avaible information about smoking behavior. As much as 70-90% of cannabis users have been reported also to smoke cigarettes^73,74^, which impose the risk of confounding. However, smoking is also highly comorbid with psychiatric disorders^75,76^, which is prevalent among the control group (66,79% was diagnosed with at least one of the major psychiatric disorders studied in iPSYCH; supplementary Table 2). Since we expect smoking to be prevalent both among cases and the control group, it is unlikely that smoking alone would generate the strong association we observe on chromosome 8. Additionally, SNPs located in or near *CHRNA3, CHRNA4, CHRNB3, CYP2A6* previously found to demonstrate extreme strong association with nicotine dependence and smoking behavior^71,72,77–79^, was not associated in the present analysis (lowest P=0.31 for rs1051730; Supplementary Table 4), and the risk locus for CUD identified in our GWAS did not demonstrate strong association with smoking behaviour in the published large GWASs of smoking^71,72,77–79^ (lowest P-value for life time smoking (P=0.003); Supplementary Table 5).

In order to evaluate the genetic overlap between CUD and a range of other phenotypes at the polygenic level, we conducted analyses of polygenic risk scores (PRS) for 22 phenotypes related to cognition, personality, psychiatric disorders, reproduction and smoking behaviour (online methods; list of phenotypes and results can be found in Supplementary Table 6). PRS for eight phenotypes (three measures of cognitive perfomance, age at first birth, life time smoking, ADHD, depressive symptoms and schizophrenia) demonstrated strong association with CUD (4.33×10^−4^ < P < 7.44×10^−15^; Figure 4 and Supplementary Table 6). Strikingly, PRS for measures of educational attainment, SSGAC^80,81^ (educational years: z-score= −7.78; P=7.44×10^−15^ and college completion z-score=-5.10; P=3.33×10^−7^), were strongly negatively associated with the risk of CUD. A finding which was reinforced by the significant negative association of PRS for human intelligence^82^ with CUD (z-score = −3.51; P=4.33×10^−4^). Numerous epidemiological studies have observed an association of cannabis use disorder and decreased educational performance^83–86^. The direction of causation is unclear and it is unknown if the relationship is caused by cannabis use and subsequent disengagement from education or wether and to what extent the two share overlapping genetic factors ^87,88^. Our results suggest an overlap in genetic risk factors, with a decrease in the odds ratio for cannabis use disorder with increased number of educational years/cognitive performance (Supplementary Figure 4.A-C.). The decreased risk of CUD with increased age at first birth (z-score =−7.41; P=1.26×10^−13^) is supporting the relationship with educational attainment as number of educational years is known to correlate genetically with later birth of the first child^89^. In order to avoid confounding in the PRS analyses of psychiatric disorders we excluded individuals among CUD cases and the control group with a diagnosis of the psychiatric disorder being analysed. PRS for ADHD^90^ (z-score =5.10; P=3.45×10^−7^), depressive symptoms^91^ (z-score 4.34; P=6.58×10^−6^) and schizophrenia^26^ (z-score=5.47; P=4.45×10^−8^) were all associated with an increased risk for CUD (Figure 4). The overlap in common genetic risk factors for cannabis use disorder with depressive symptoms is in line with recent observations^92^ and the association of PRS for ADHD with CUD suggest that the epidemiological observations of high comorbidity of ADHD with CUD^93,94^ are influenced by shared genetic risk components. Likewise, the PRS analysis supports an overlap in common genetic risk factors between CUD and schizophrenia (in line with epidemiological observations^33,34^). However we cannot rule out if the overlap involves variants with pleiotropic effect on the two disorders, or if the overlap in general is caused by heterogeneity among schizophrenia cases due to a subgroup comorbid with CUD, in line with the CUD risk locus identified here, which does not seem to have a pleitropic effect across the two disorders.

In summary, we identified a genome-wide significant CUD locus on chromosome 8, which replicated in an independent sample. An impact of the locus on earlier age of first diagnosis was further suggested. The locus implicates *CHRNA2* in CUD through analyses of the imputed genetically regulated gene expression, showing decreased *CHRNA2* expression in cerebellum (and other brain regions) of individuals with cannabis use disorder. An intriguing result which suggests that CHRNA2 could be a potential therapeutic target for further testing in randomized controlled trials. Analyses of PRS for other phenotypes revealed a significant decrease in the risk of CUD with increased measures of cognitive performance.

## Online Methods

### Sample

The individuals in this study is a part of the large Danish nation-wide population based sample collected by iPSYCH^25^. The iPSYCH sample was originally designed for studying major psychiatric disorders and includes genotypes from 79,492 individuals, including 54,249 cases diagnosed with at least one of six mental disorders (schizophrenia, bipolar disorder, major depressive disorder, ADHD, anorexia and autism spectrum disorder) and 25,243 individuals not having any of the six psychiatric disorders. The iPSYCH sample was selected from a birth cohort comprising individuals born in Denmark between May 1, 1981, and December 31, 2005, who were residents in Denmark on their first birthday and who have a known mother (N = 1,472,762). The iPSYCH cases were identified based on information in the Danish Psychiatric Central Research Register^95^, and 30,000 randomly selected controls were identified from the same nationwide birth cohort in the Danish Civil Registration System^96^. Subseqeuntly blood spot samples (Guthrie cards) were identified in the in the Danish Newborn Screening Biobank (DNSB) from the included individuals^97^. Processing of DNA, genotyping and genotype calling as well as imputing of genotypes (see below) of the samples were done in 23 waves of approximately 3,500 individuals each.

Individuals with CUD were identified in the iPSYCH sample through information in the Danish Psychiatric Central Research Register^95^ as well as in the Danish National Patient Register^98^ using information up to 2013. All individuals with an ICD10 diagnosis of F12.1-12.9, were included. Individuals with acute intoxication (diagnosis code F12.0) were not included. The F12.0 diagnosis is related to the acute pharmacological effects of cannabis use and resolve with time, and does not necessary reflect long term problematic cannabis use, which is the focus of this study. All individuals in the iPSYCH sample not having a diagnosis of CUD were used as controls and covariates for the major psychiatric disorders studied by iPSYCH were used in the analyses in order to correct for the presence of mental disorders. Only 54 individuals had a diagnosis of CUD and anorexia, and all individuals having a diagnosis of anorexia among cases and the control group were excluded. Supplementary Table 2 gives an overview of the distribution of individuals with a psychiatric disorder among the CUD cases and the control group.

Processing of samples, genotyping, QC and imputing was done in 23 waves. In this study individuals from the first four waves were excluded as only a few cannabis use disorder cases were present in these waves (range 0-15 cases). The waves represent approximate birth years and the four waves that were excluded represent to a large extend individuals, which in 2013 (the time of extraction of register diagnoses) were children.

### Genotyping, quality control and GWAS

DNA was extracted from dried blood spot samples and whole genome amplified in triplicates as described previously^99,100^. Genotyping was performed at the Broad Institute of Harvard and MIT (Cambridge, MA, USA) using Illumina’s Beadarrays (PsychChip; Illumina, CA, San Diego, USA) according to the manufacturer’s protocols. Genotypes were a result of merging call sets from three different calling algorithms (GenCall, Birdseed and Zcall). GenCall^101^ and Birdseed^102^ was used to call genotypes with minor allele frequency (maf) > 0.01 and zCall^103^ was used to call genotypes with maf < 0.01. The merging was done after pre-QC on individual call sets.

Stringent quality control was applied to the individual call rate (> 0.98) and only genotypes with high call rate (> 0.98), no strong deviation from Hardy-Weinberg equilibrium (*P* > 10^−6^ in controls or *P* > 10^−10^ in cases) and low heterozygosity rates (| F_het_ | < 0.2) were included. Genotypes were phased and imputed using the 1000 Genomes Project phase 3 (1KGP3)^29,104^ imputation reference panel and SHAPEIT^27^ and IMPUTE2^28^. Relatedness and population stratification were evaluated using a set of high quality markers (genotyped autosomal markers with minor allele frequency (maf) > 0.05, HWE P > 1 × 10^−4^ and SNP call rate > 0.98), which were pruned for linkage disequilibrium (LD) (r^2^ < 0.075) resulting in a set of 37,425 pruned markers (markers located in long-range LD regions defined by Price et al.^105^ were excluded). Genetic relatedness was estimated using PLINK v1.9^106,107^ to identify first and second-degree relatives 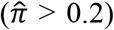 and one individual was excluded from each related pair (cases preferred kept over controls). Genetic outliers were identified for exclusion based on principal component analyses (PCA) using EIGENSOFT^108,109^. A genetic homogenous sample was defined based on a subsample of individuals being Danes for three generations (identified based on register information about birth country of the individuals, their parents and grandparents). The subsample of Danes was used to define the center based on the mean values of principal component (PC) 1 and PC2. Subsequently PC1 and PC2 were used to define a genetic homogenous population by excluding individuals outside an ellipsoid with axes greater than six standard deviations from the mean.

Association analysis was done using logistic regression and the imputed marker dosages. The following covariates were used: principal component 1-4 and principal components from the PCA associated with case-control status, the 19 data-processing waves and diagnosis of major psychiatric disorders studied by iPSYCH (supplementary Table 2). Results for 9,729,295 markers were generated, subsequently markers with imputation info score < 0.7 (n = 608.367), markers with maf < 0.01 (n=10.220) and multi-allelic markers (n = 143,083) were removed. In total after filtering 8,969,939 markers remained for further analysis. All analyses of the iPSYCH sample were performed at the secured national GenomeDK high performance-computing cluster in Denmark (<u>https://genome.au.dk</u>).

### Replication

The genome-wide significant locus on chromosome 8 was replicated in an independent European cohort consisting of 5,501 cases with diagnosed CUD and 301,041 population controls collected by deCODE genetics. The characteristics of the SAA treatment sample have been described previously^110^ and currently diagnoses at Vogur Hospital are made using DSM-V, but most of the diagnoses in this study are based on DSM-IV or DSM-IIIR. The genotypes were obtained based on SNP array data and whole genome sequences using long range phasing and two types of imputations^111^. For the replication study, markers were looked up in the results of a genome-wide association study performed by logistic regression treating disease status as the response and genotype counts as covariates. Other available individual characteristics that correlate with disease status were also included in the model as nuisance variables, using previously described methods^112^. The resulting P values were corrected for inflation by the method of genomic controls (correction factor=1.42). Nine genetic variants all representing the same association signal on chromosome 8 with P values less than 1×10^−6^ were looked up in the deCODE results. The variants in the locus were selected based on LD (0.2 < r^2^ < 0.7). We included additional markers besides the index SNP in order to be able evaluate the consistency of direction of association in the replication cohort over a set of markers located in the associated risk locus. The data were meta-analyzed using summary statistics and inverse variance weighted fixed effects model, implemented in the software METAL^113^. All tested variants demonstrated consistent direction of association in the replication cohort.

### Heritability

SNP heritability (h^2^_SNP_) was estimated by LD score regression^30^ using summary statistics from the GWAS of CUD and pre-computed LD scores (available on https://github.com/bulik/ldsc). The SNP heritability was calculated on the liability scale using a prevalence of 1% in the population. The h^2^_SNP_ estimated by LD score regression was inaccurate as the ratio between the estimated h^2^_SNP_ and the standard error was less than four (h^2^_SNP_/SE = 2,86). Therefore, the h^2^_SNP_ was also estimated using GCTA^31^. A genomic relationship matrix (GRM) between all pairwise combinations of individuals was estimated for each autosome separately and subsequently merged using the GCTA software. The strict best-guess-genotypes (i.e. 4,299,887 SNPs with info score > 0.8, missing rate < 0.01 and MAF > 0.01) from imputation with phase 3 of 1000 genomes were used for GRM estimation. Univariate GREML analyses in GCTA was used to estimate h^2^_SNP_ on the liability scale, using the combined GRM and the same covariates as in the GWAS.

### Leave-one phenotype out GWAS

In the GWAS of CUD we corrected, by covariates, for five major psychiatric disorders analysed in iPSYCH (Supplementary Table 2). However, in order to further evaluate the potential impact on the association signal of the index SNP from the psychiatric phenotypes, sensitivity analyses were performed where one phenotype at a time was excluded from the association analysis. The phenotypes evaluated were schizophrenia, bipolar disorder, major depressive disorder, attention-deficit hyper activity disorder, autism and individuals not having any of the five diagnoses. GWASs were performed as previously described excluding one of the six phenotypes one at a time (sample size for number of cases and controls is shown in Table 2).

### Permutation based evaluation of the odds ratio of rs56372821 for schizophrenia with and without CUD

We applied a permutation based approach in order to evaluate how the inclusion and exclusion of individuals with CUD affected the association of our index SNP rs56372821 with schizophrenia. The analyses were based on the iPSYCH sample consisting of 2,281 schizophrenia cases and 23,134 population based controls (genotyping, QC and imputation were done using the same procedures as explained above). Odds ratios were obtained by the glm() function in R using dosage data from the imputed SNP rs56372821, and including principle components PC1-PC4 + PCs signicantly associated with SZ and/or CUD and wave number as covariates.

In the association analysis of schizophrenia the odds ratio of 56372821(CI95%) was OR=0.9 (0.8-10.99). When excluding all individuals with CUD, among cases and controls (1,727 cases and 23,033 controls) the odds ratio of rs56372821(CI95%) was OR = 0.97 (0.87-1.09). To evaluate whether this change in the odds ratio is due to removal of individuals with CUD (554 cases and 101 control with CUD removed) or due to reduction in sample size, a null distribution of OR’s was generated by randomly removing 554 individuals among schizophrenia cases and 101 individuals among controls (removing a total of 655 individuals). This permutation of sample exlcusion was done 9,999 times to generate the null distribution of the odds ratio for rs56372821. To obtain a two-sided P-value, the observed ln(OR_rs56372821_) = −0.03 was mirrored around the mean ln(OR_rs56372821_) of the null distribution, as shown in Supplementary Figure 1. Removal of individuals with CUD (554 SZ cases and 101 controls) changed the OR of rs56372821 significantly, compared to random removal of the same number of schizophrina cases and controls (P_two-sided_=0.0027), producing a significant increase of OR_rs56372821_ compared to random removal (P_one-sided_=0.0015).

### Association with age at first diagnosis

In order to test for a potential impact of the risk locus on age at first diagnosis, a case only study was performed testing for association of the index SNP (rs56372821) and SNPs in LD with this (r^2^ > 0.7; 19 variants) with age at first diagnosis. Date of diagnosis was identified from register information in the Danish Psychiatric Central Research Register^95^. Analysis for association with age at first diagnosis (natural logarithm (age at first diagnosis)) was done using linear regression and the same covariates as used in the GWAS (PCs, wave and the psychiatric disorders). A dominant model with respect to the risk allele, was applied.

### PrediXcan

The genetically regulated gene expression was imputed in 11 brain tissues using PrediXcan^35^ (models downloaded from: <u>https://github.com/hakyimlab/PrediXcan</u>; version 6 data). PrediXcan was used to impute the transcriptome for cases and the control group using SNP weights derived from models trained on reference transcriptome data sets including 10 brain tissues from GTEx (https://www.gtexportal.org/home/) and transcriptome data from dorsolateral prefrontal cortex generated by the CommonMind Consortium^37,(Huckins et al. submitted to Nature Genetics)^. The models were trained on 1000 genome SNPs and contained only gene expression prediction models with a false discovery rate less than 5%. Gene expression levels in the iPSYCH data were imputed wave-wise and subseqeuntly the imputed data were merged. We tested for association of 2,459 – 10,930 protein coding genes (depending on the tissue; see Supplementary Table 3), with CUD using logistic regression corrected by the relevant principal components from PCA and the psychiatric disorder as described for the GWAS. Since gene expression among the different brain tissues is highly correlated, evaluated based on GTEx data^114^, we corrected the P-value for each gene by the total number genes tested in all tissues with a valid model available for the gene, with respect to *CHRNA2* we corrected for 13,166 genes tested in three tissues.

### PRS risk score analyses

PRS analyses were done using GWAS summary statistics from 22 GWASs (Supplementary Table 6). The summary files were downloaded from public databases and processed using the munge script which is a part of the LDscore regression software^30^. All variants with INFO < 0.9, MAF < 0.01, missing values, out of bounds P-values, ambiguous strand alleles and duplicated rs-ids were removed using the munge script. In addition, mult-allelic variants and insertion and deletion (indels) were removed. The processed summary files were then LD-clumped using Plink, with the following parameter settings: –clump-p1 1 –clump-p2 1 –clump-r2 0.1 –clump-kb 500. The clumped file were used as the training dataset. Genetic risk scores were estimated at different P-value thresholds for SNP inclusion: 5×10^−8^, 1×10^−6^, 1×10^−4^, 1×10^−3^, 0.01, 0.05, 0.1, 0.2, 0.5 and 1.0 for all individuals in the target sample (CUD cases and the control group) from the genotype dosages using Plink’s ‘–score’ method, with default arguments. However the PRS scores for ADHD, were generated using the approach described Demontis et al.^90^. For each P-value threshold the variance in the phenotype explained by PRS was estimated using Nagelkerke’s *R* (R package ‘BaylorEdPsych’)^*2*^, and association of PRS with CUD was estimated using logistic regression including the same covariates used in the GWAS analysis (PCs from PCA and the psychiatric disorders listed in Supplementary Table 2). In PRS analyses of psychiatric disorders (ADHD, schizophrenia and depression related phenotypes) individuals with a diagnosis of the disorder being analysed were excluded. The number of individuals excluded with ADHD, schizophrenia and major depressive disorder are listed in Supplementary Table 2.

## Acknowledgements

We are grateful to the following researchers for providing the results for the replication: Thorgeir E. Thorgeirsson^1^, Gunnar W. Reginsson^1^, Thorarinn Tyrfingsson^2^, Valgerdur Runarsdottir^2^, Hreinn Stefansson^1^, Kari Stefansson^1^ (^1^deCODE genetics / Amgen, Reykjavik, Iceland; ^2^National Center of Addiction Medicine, SAA, Vogur Hospital, Reykjavik, Iceland)

The iPSYCH project is funded by the Lundbeck Foundation (grant numbers R102-A9118 and R155-2014-1724) and the universities and university hospitals of Aarhus and Copenhagen. Genotyping of iPSYCH samples was supported by grants from the Lundbeck Foundation, the Stanley Foundation, the Simons Foundation (SFARI 311789 to MJD), and NIMH (5U01MH094432-02 to MJD). The Danish National Biobank resource was supported by the Novo Nordisk Foundation. Data handling and analysis on the GenomeDK HPC facility was supported by NIMH (1U01MH109514-01 to Michael O’Donovan and ADB). High-performance computer capacity for handling and statistical analysis of iPSYCH data on the GenomeDK HPC facility was provided by the Centre for Integrative Sequencing, iSEQ, Aarhus University, Denmark (grant to ADB).

We gratefully acknowledge all the studies and databases that made GWAS summary data available: GPC (Genetics of Personality Consortium), PGC (Psychiatric Genomics Consortium), SSGAC (Social Science Genetics Association Consortium), TAG (Tobacco and Genetics Consortium), UK Biobank, CTGLAB (Complex Traits Genetics Lab).

We gratefully acknowledge the data made available by the the Genotype-Tissue Expression (GTEx) Project (supported by the Common Fund of the Office of the Director of the National Institutes of Health, and by NCI, NHGRI, NHLBI, NIDA, NIMH, and NINDS). The data used for the analyses described in this manuscript were obtained from: https://www.gtexportal.org/home/ on 01/11/2017.

## Disclosures

T. Werge has been a lecturer and advisor to H. Lundbeck A/S.

T. E. Thorgeirsson, G. W. Reginsson, H. Stefansson and K. Stefansson are employees of deCODE genetics/Amgen.

## Author contributions

This section will be filled during the review process

